# Phylogenomics of plant-associated *Botryosphaeriaceae* species

**DOI:** 10.1101/2021.01.12.426103

**Authors:** Jadran Garcia, Daniel P. Lawrence, Abraham Morales-Cruz, Renaud Travadon, Andrea Minio, Rufina Hernandez-Martinez, Philippe E. Rolshausen, Kendra Baumgartner, Dario Cantu

## Abstract

The *Botryosphaeriaceae* is a fungal family that includes many destructive vascular pathogens of woody plants (e.g., Botryosphaeria dieback of grape, Panicle blight of pistachio). Species in the genera *Botryosphaeria, Diplodia, Dothiorella, Lasiodiplodia, Neofusicoccum*, and *Neoscytalidium* attack a range of horticultural crops, but they vary in virulence and in their abilities to infect their hosts via different infection courts (flowers, green shoots, woody twigs). Isolates of seventeen species, originating from symptomatic apricot, grape, pistachio, and walnut were tested for pathogenicity to grapevine wood after four months of incubation in potted plants in the greenhouse. Results revealed significant variation in virulence in terms of the length of the internal wood lesions caused by these seventeen species. Phylogenomic comparisons of the seventeen species of wood-colonizing fungi revealed clade-specific expansion of gene families representing putative virulence factors involved in toxin production and mobilization, wood degradation, and nutrient uptake. Statistical analyses of the evolution of the size of gene families revealed expansions of secondary metabolism and transporter gene families in *Lasiodiplodia* and of secreted cell wall degrading enzymes (CAZymes) in *Botryosphaeria* and *Neofusicoccum* genomes. In contrast, *Diplodia, Dothiorella*, and *Neoscytalidium* generally showed a contraction in the number of members of these gene families. Overall, species with expansions of gene families, such as secreted CAZymes, secondary metabolism, and transporters, were the most virulent (i.e., were associated with the largest lesions), based on our pathogenicity tests and published reports. This study represents the first comparative phylogenomic investigation into the evolution of possible virulence factors from diverse, cosmopolitan members of the *Botryosphaeriaceae*.

## Introduction

The fungal family *Botryosphaeriaceae* (*Botryosphaeriales, Dothideomycetes*) was introduced in 1918 by Theissen & Sydow with *Botryosphaeria* as the type genus. Members of this group have been taxonomically characterized based on the production of large, ovoid to oblong, typically hyaline, aseptate ascospores, which may become brown and septate with age, within bitunicate asci within unilocular or multilocular botryose ascomata known as pseudothecia (Sivanesan 1984; Phillips *et al*., 2005). The asexual states of *Botryosphaeriaceae* exhibit a wide range of conidial morphologies that are taxonomically informative (Phillips et al., 2005). Crous et al. (2006) contributed to stabilize the taxonomy of the genera within the *Botryosphaeriaceae* by employing a natural unit classification scheme, which is also referred to as the “genus-for-genus concept” (Seifert et al., 2000). The distinct asexual morphs were linked to unique sexual morphs on a unit-by-unit basis, which was corroborated with phylogenetic analysis of 28S rDNA sequence data revealing 10 generic clades. The *Botryosphaeriaceae* is currently composed of 24 well-defined genera and more than 200 species (Burgess et al., 2019) that are cosmopolitan in distribution and exist primarily as saprobes, endophytes, or pathogens on a wide array of important perennial plant hosts (Slippers & Wingfield 2007), in both human-altered (agricultural and urban) and natural ecosystems (forests and riparian areas) (Slippers et al., 2009; Lawrence et al., 2017).

The ecology of Botryosphaeriaceous taxa is complex and not fully understood. For example, in spite of being a shoot blight and canker pathogen of pine, *Diplodia sapinea* has been isolated from the bark surface and internal woody tissues of woody twigs from asymptomatic *Pinus* (Petrini & Fisher 1988), representing what some may consider an ‘endophytic phase’, in which neither the internal plant tissues from which it is isolated, nor other plant tissues/organs showed apparent symptoms, nor were there negative impacts to host growth at the time of isolation. A similar pattern in the ecology of other *Botryosphaeriaceae* species considered pathogenic, but later being isolated during an endophytic phase, has been documented (Slippers & Wingfield 2007, Luo et al., 2019, Hrycan et al., 2020). In some cases, abiotic stress (water stress, heat stress) has been shown to induce severe symptoms in different host plants infected with seemingly innocuous *Botryosphaeriaceae* (Pusey 1989; Mullen et al., 1991; Smith et al., 1994). This relationship between abiotic stress and more severe symptoms or more rapid colonization has also been reported for pathogenic species, e.g., *Neofusiococcum parvum* causing Botryosphaeria dieback of grape (Luque et al., 2010; Galarneau et al., 2019) and *Botryosphaeria dothidea* causing Pistachio panicle and shoot blight (Ma et al., 2001). Under climate-change scenarios of more frequent temperature extremes and prolonged drought, the interactions between host plants and Botryosphaeriaceae species may transit more readily from endophytic to pathogenic (Desprez-Loustau et al., 2006; Slippers et al., 2007). An increase in *Botryosphaeriaceae* symptom severity in conjunction with other biotic stresses has also been documented in the literature (Lawrence et al., 2018, Old et al., 1990).

Members of the *Botryosphaeriaceae* are probably most well-known as being destructive blight and canker pathogens of planted hosts (Luo et al., 2019). In agricultural settings, for example, they infect a large number of fruit and nut crops, such as almond (Inderbitzin et al., 2010; Gramaje et al., 2012; Nouri et al., 2018; Holland et al., 2020), apple (Phillips et al., 2012), avocado (McDonald et al., 2009), citrus (Linaldeddu et al., 2015), grapevine (Úrbez-Torres 2011), olive (Úrbez-Torres et al., 2013), pistachio (Michailides 1991; Nouri et al., 2019), and walnut (Chen et al., 2014a). In forest plantations in Australia and South Africa, for example, they infect *Eucalyptus* spp. and *Pinus* spp. (Slippers et al., 2007). Infection is through either wounds to green and woody tissues or through natural openings in flowers, fruit, leaves, and shoots. The pathogens produce enzymes and/or toxins that kill cells and tissues of the various plant organs they attack. Infections of woody tissues of perennial hosts, either deep in the wood or just below the bark, can lead to stunted shoot growth, with eventual shoot death or ‘dieback’.

Ecological genomic comparisons of phytopathogenic and saprobic fungi suggest that the former possess expanded gene families that generally fall into two main functional categories: 1) lytic capabilities (Massonnet et al., 2018) and 2) putative transporters (Powell et al., 2008). Fungal lignin peroxidases, peroxidases, laccases, and polyphenol oxidase allow fungi to gain access to nutrients and to protect themselves from host defenses while growing in wood (Martínková et al. 2016; Mayer 2006; Valette et al. 2017). Pathogenic species with the ability to enzymatically decompose a broader diversity of cell wall carbohydrates might be expected to more rapidly colonize, kill, and/or decompose host tissue. Membrane transporters of fungal plant pathogens also play important roles in exporting virulence factors involved in pathogenesis, influx of nutrients, and efflux of host-derived defense antimicrobial compounds (Denny and VanEtten 1983; Denny et al. 1987). Previous genomic comparisons of phylogenetically diverse wood-infecting pathogens of grape revealed expansions in the repertoire of cell-wall degrading enzymes called carbohydrate-active enzyme (CAZyme) gene families, whose protein products are involved in the creation, degradation, and/or modification of glycosidic bonds of plant cell wall constituents, including the main components of wood, cellulose, hemicelluloses, lignin (Morales-Cruz et al., 2015), and significantly so in *Neofusicoccum parvum*. Further, a recent genomic annotation and *in planta* transcriptomic study of putative virulence factors of *Neof. parvum* during wood colonization revealed 567 protein-coding genes belonging to 52 different CAZyme families with glycoside hydrolases (GHs), which made up approximately 50% of the pathogen’s cell-wall degrading repertoire (Massonnet et al., 2018). Likewise, Yan et al. (2018) identified 820 CAZymes with at least 10 families that have experienced expansion in the genome of *L. theobromae* with GHs representing the largest super family involved in the modification of plant cell wall carbohydrates. Genome comparisons of *Botryosphaeria dothidea, L. theobromae*, and *Neof. parvum* revealed that the genome of *L. theobromae*, the most virulent of the three species, is expanded in gene families associated with membrane transport, mainly ATP-binding-cassette (ABC family) and major facilitator super (MFS) families (Yan et al., 2018). That same study reported 17 membrane transport genes that were significantly up-regulated upon host recognition including amino acid transporters and sugar porters. The largest transporter families reported in the genome of *Neof. parvum* include MFS, Peroxisomal Protein Importer (PPI) family, and the ABC superfamily (Massonnet et al., 2018).

In this study we analyze the genome sequences of seventeen *Botryosphaeriaceae* species representing six genera (*Botryosphaeria, Diplodia, Dothiorella, Lasiodiplodia, Neofusicoccum*, and *Neoscytalidium*), which are wood-canker pathogens that attack horticultural crops, namely grape, pistachio, *Prunus* species (almond and stone fruits apricot, peach, and plum), and walnut. Our objective is to examine through phylogenomic comparisons this comprehensive set of species on one host, grape, to better understand the evolutionary trends within this important fungal family, especially as it pertains to the gene space involving pathogenesis of woody tissues and fungal virulence.

## Materials and Methods

### Isolate collection and species confirmation

All fungal isolates utilized in this study were obtained from internal wood cankers of symptomatic hosts following the protocol of Baumgartner et al. (2013) (**Table 1**). Total genomic DNA was extracted following Morales-Cruz et al. (2018). The internal transcribed spacer (ITS) and translation elongation factor (TEF) loci were amplified for each isolate via PCR using primers ITS5/ITS4 (White et al., 1990) and EF1-688F/EF1-1251R (Alves et al., 2008). TEF and ITS sequences of each species (including type specimen sequences downloaded from GenBank) were concatenated and aligned using MUSCLE v3.8.31 (121) with default parameters. The alignment was cleaned with GBlocks v. 0.91b (Castresana, 2000) with a minimum block’s length of 5 bp and half of the gaps allowed. PhyML (Guindon *et al*., 2010) was used to calculate the maximum likelihood tree using 100 bootstrap replications, HKY85 substitution model and the subtree-pruning-regrafting method for searching for optimal tree topology. The resulting tree was visualized and edited for presentation using FigTree v1.4.1 (Rambaut 2012).

**Table 1:**
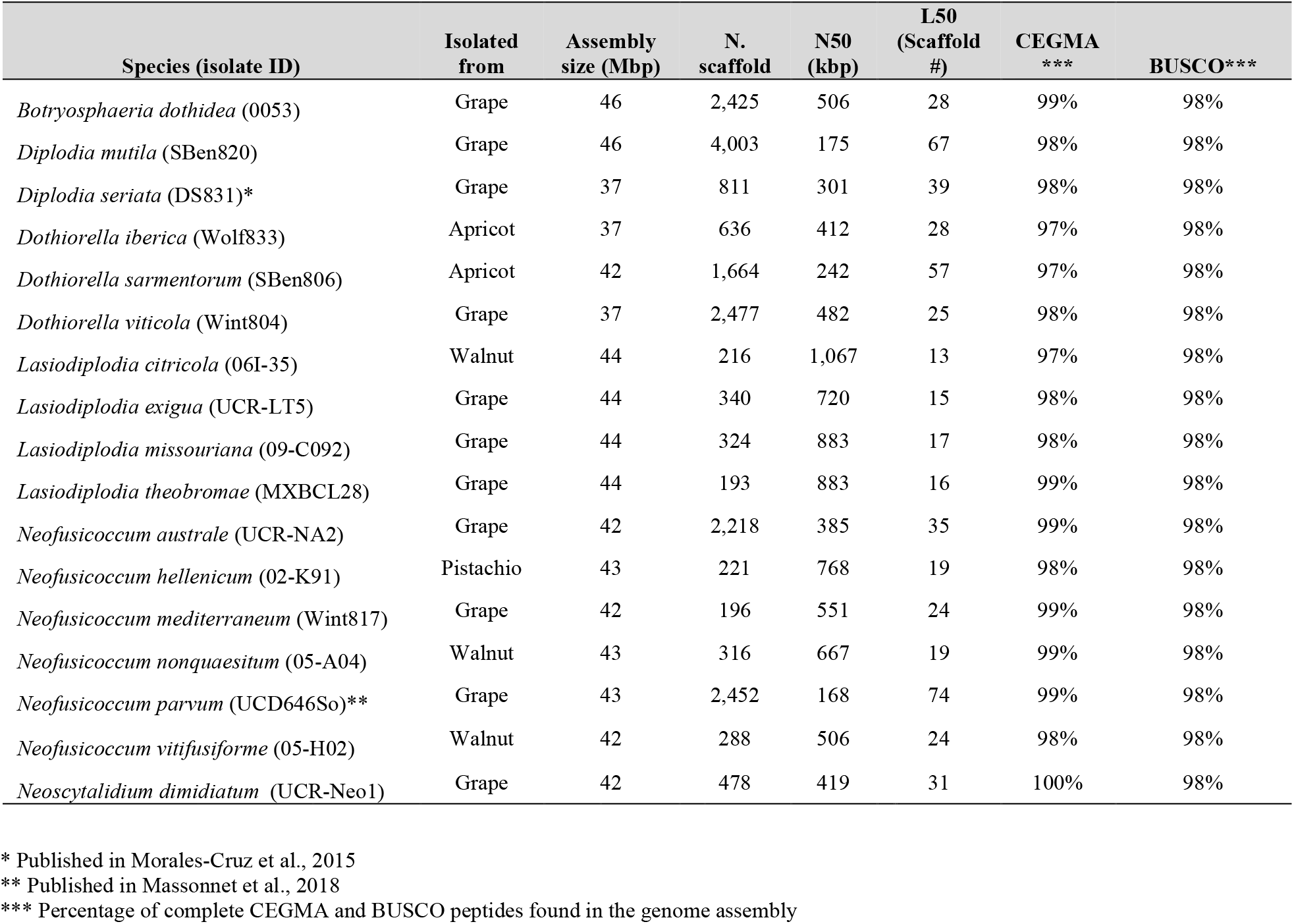
Genome assembly summary statistics of the *Botryosphaeriaceae* species analyzed.

### Sequencing and Genomes assemblies

DNA extraction was done following the methods used by Morales-Cruz et al. (2015) using the axenic cultures of the isolated fungi and a CTAB protocol. Sequencing libraries were prepared and sequenced as described in Morales-Cruz et al. (2015). Raw reads were trimmed for quality (Q>30) and adapter removal using Trimmomatic v0.36 (Bolger *et al*., 2014). Assembly of high-quality reads was made using SPAdes v3.9 (Bankevich *et al*., 2012) with the careful option and automatic read coverage cutoff. Assembly completeness was assessed using the Core Eukaryotic Genes Mapping Approach (CEGMA v.2.5; Parra *et al*., 2007) and Benchmarking Universal Single-Copy Orthologs (BUSCO v.1.1; Simão *et al*., 2015) analysis. RepeatMasker v.4.06 (Smit *et al*., 1996-2015) with default parameter was used to mask repeats. Gene model prediction was performed with Augustus v.3.2.1 (Stanke *et al*., 2006) with default parameter and using *Neofusicoccum parvum* gene model as training set. Sequencing data are available at NCBI (Bioproject PRJNA673527). Sequencing data of *Di. seriata* (Morales-Cruz et al., 2015) and *Neof. parvum* (Massonnet et al., 2018) can be retrieved from NCBI under Bioproject PRJNA261773 and PRJNA321421, respectively. All genome assemblies and gene models are publicly available at Zenodo (DOI: 10.5281/zenodo.4417445).

### Functional annotation

The general annotation of the predicted proteins was assigned based on the similarities with peptides in the GenBank with Blast2GO (Conesa *et al*., 2005), and to conserved domains in Pfam database (Finn *et al*., 2016). The functional annotation (**Additional file 1 Table 3**) was assigned based on the databases and parameters presented in **Additional file 1 Table 4**. CAZymes were annotated with the dbCAN2 (Zhang *et al*., 2018). The signal peptides were assigned using SignalP 5.0 (Armenteros *et al*., 2019). The proteins with annotation in both databases were annotated as secreted CAZymes. Secondary metabolites clusters were annotated using antiSMASH 5.0 (Blin *et al*., 2019). Peroxidases were annotated using a specialized database for fungi called fPoxDB (Choi *et al*., 2014). CYPED 6.0 was used to annotate the Cytochrome P450 proteins (Fischer *et al*., 2007). Last, the proteins involved in transportation functions were annotated using the TCDB (Saier Jr *et al*., 2006; Saier Jr *et al*., 2016).

### Construction of a clock-calibrated phylogenetic tree

Seventy-three single copy peptides used in (Floudas et al., 2012) for fungi phylogeny reconstruction were extracted from the reference strain *Saccharomyces cerevisiae* 2S88C Genome Release 64-2-1 (downloaded from http://www.yeastgenome.org). All these peptides were compared using BLASTP (v.2.6.0+) against the seventeen *Botryosphaeriaceae* species and two wood-decay basidiomycetes that colonize grape: pathogenic, wood-rotting fungus (with characteristics of both white-rot and brown-rot fungi) *Fomitiporia mediterranea* and saprobic, white-rot fungus *Stereum hirsutum. F. mediterranea* is one of a complex of pathogens that causes the grapevine trunk disease Esca in Europe, whereas the pathogenicity of *S. hirsutum* to grape is not known (Fischer 2006). Twenty-one proteins had exactly one top hit in all the species. The rest of the seventy-three initial proteins were excluded because they were not present in all the species or because they had paralogs. Each set of orthologous proteins was aligned using MUSCLE v3.8.31 (Edgar, 2004). Alignments were concatenated and cleaned using GBlocks v. 0.91b (Castresana, 2000; maximum number of contiguous nonconserved positions= 4, minimum length of a block= 10), reducing the initial 21008 positions to 12066 really informative positions. Clean alignments were imported into BEAUti (v1.10.4) to prepare them for BEAST (v1.10.4) analysis (Bouckaert et al., 2014). Monophyletic partitions were set for *Ascomycota, Basidiomycota* and *Dothideomycetes* species. Calibrations points were set to 588 and 350 mya on *Ascomycota* and *Dothideomycetes* partition, respectively according to (Beimforde *et al*., 2014). Six MCMC chains of 1,000,000 steps were launched on BEAST (WAG substitution model, 4 Gamma Categories + Invariant sites, Lognormal relaxed clock, Calibrated Yule Model). The resulting trees were concatenated with LogCombiner (v1.10.4; Bouckaert et al., 2014) and a consensus tree was obtained from TreeAnnotator (v1.10.4; (Bouckaert et al., 2014).) (**Additional file 2**). FigTree v1.4.1 (Rambaut, 2012) and Inkscape v.1.0.1 (Inkscape Project, 2020) were used to edit the tree for figure presentation.

### Computational analysis of gene family evolution (CAFE)

BLASTP (e-value < 10^−6^) as used to group proteins in families based on sequence similarity followed by Markov clustering with MCL (Van Dongen 2000; Enright *et al*., 2002). 10158 families with at least one protein in no less than four species were used with the clock calibrated tree as input for the CAFÉ v.4.2.1 (De Bie *et al*., 2006) analysis. CAFE was run in default mode with the option -s to optimize the lambda parameter to 0.00155948837239, and a P-value threshold of 0.01 (option -p). To evaluate significant expansion or contraction of a specific branch, Viterbi P-values were calculated for each significant family.

### Phylogenetic principal component analysis

The phyl.pca function of the phytools R package (Revell 2012) was used to create the phylogenetic PCAs. The inputs for his function were the clock calibrated tree and protein matrices of CAZymes and Secondary metabolites clusters.

### Pathogenicity tests

The pathogenicity of 17 species of *Botryosphaeriaceae* was evaluated on potted grapevines (Pinot Noir clone 777) in replicate experiments in the greenhouse (18 treatments × 10 replicate plants per treatment × 2 experiments = 360 total plants). Hardwood cuttings obtained from a commercial nursery were callused, inoculated, and then rooted in pots in May/June 2016 following the protocol of Travadon et al. (2013). For inoculations, a power drill was used to wound (5 mm wide × 3 mm deep) the cutting, approximately 2 cm below the apical node. A 5-mm agar plug from a 7-day culture on PDA was aseptically inserted into the wound and sealed with Vaseline and parafilm to prevent inoculum desiccation. Cuttings were coated in melted paraffin wax (Gulf Wax; Royal Oak Enterprises, LLC, Roswell, GA) and potted in a sterile potting mix amended with slow-release fertilizer (Osmocote® Pro 24-4-9, Scotts, Marysville, OH). The plants were watered twice per week for 16 weeks. Ten plants were used for each isolate and ten plants were mock-inoculated with sterile PDA. Plants were arranged in a completely randomized design in two separate greenhouses at the University of California Experiment Station in Davis from June 2016-October 2016 (natural sunlight photoperiod, 25 +/- 1 C [day], 18 +/- 3 C [night]. The second experiment was initiated one week after the first experiment. The length of internal wood discoloration extending out from the inoculation site up and down the stem (lesion length) was measured approximately 16 weeks after inoculation in October 2016. First, plants were inspected for foliar symptoms. Then the newly developed green shoots, roots, and bark of each plant were removed and discarded, and the woody stems were surface sterilized in 1% sodium hypochlorite for 2 min and rinsed with deionized water. The length of each stem was recorded and cut longitudinally to expose wood discoloration, the length of which was measured with a digital caliper.

Lesion lengths were used as a measure of pathogenicity. Normality and homogeneity of variances were evaluated using Shapiro-Wilk’s and Levene’s tests, respectively. ANOVA was used to determine whether there were differences in lesion length among treatments. ANOVA was performed in R using the lesion size as a function of the inoculation treatment and the experiment. Means were compared for significant effects (P < 0.05) by Tukey’s HSD post-hoc test.

## Results

### Genome assembly, gene prediction, and virulence factor-focused functional annotation

To expand the genomic information for *Botryosphaeriaceae*, we *de novo* assembled the genomes of fifteen species isolated from multiple hosts. We included in the comparative genomics analysis the previously published genomes of *Neof. parvum* and *Di. seriata* (**Table 1** and **Additional file 1 Table 1**). For all seventeen species, pathogenicity was evaluated using inoculations of potted grapevines (**Figure 1**). All seventeen species produced dark necrotic lesions in the woody stems extending upward and downward from the point of inoculation at fifteen weeks post inoculation. Overall, *Lasiodiplodia* and *Neofusicoccum* spp. were the most aggressive, while *Diplodia* and *Dothiorella* spp. caused the smallest lesions.

**Figure 1.**
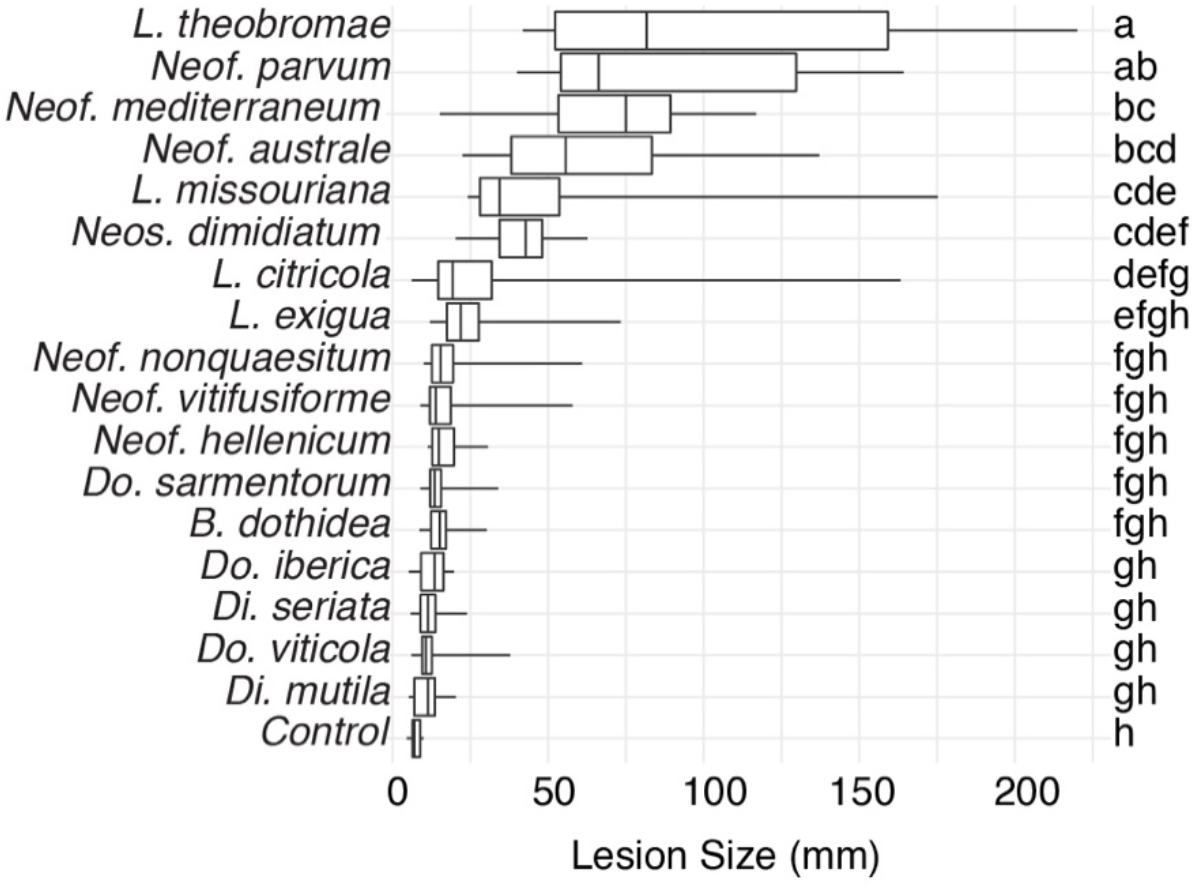
Box plots illustrating the distribution of lesion sizes distribution caused by the individual isolates of the 17 species of the *Botryosphaeriaceae* family after 4 months incubation in potted grapevine plants grown under greenhouse conditions. Bars with the same letter were not significantly different at *P* < 0.05 according to Tukey’s test.

All genomes were sequenced using Illumina technology at coverage 171 ± 10x (**Additional File 1 Table 1**). On average, sequencing Illumina reads were assembled into 1,066 ± 308 scaffolds (N50 length: 577.64 ± 64.53 kbp; L50 scaffold count: 27.9 ± 4 scaffolds). The total genome assembly size varied from 37 Mbp for *Do. viticola* to 46 Mbp for *B. dothidea* with an average of 42.37 ± 0.69 Mbp. The expected and assembled genome size had a discrepancy of less than ten percent on average (4.4 ± 2.2 Mb), which suggests near completeness of the assemblies (**Additional File 1 Table 1**). This result was confirmed by CEGMA (Parra *et al*., 2007) and BUSCOs (Simão *et al*., 2015) analyses, which reported an average 98.3 ± 0.2% and 98.1 ± 0.1% of completeness, respectively (**Table 1**). Interspersed repeats only accounted for 1.87 ± 0.003% of the genome assemblies. Among the classified elements, long-terminal-repeats (LTR) were the most abundant ranging from a total of 315 kbp in *B. dothidea* to 26 kbp in *Neos. dimidiatum* (**Additional file 1 Table 2**). The predicted protein-coding genes in the seventeen genomes varied from 10,827 in *Do. iberica* to 13,492 in *Do. sarmentorum*. On average 12,193 ± 193 CDS were found per species (**Table 2**).

**Table 2.**
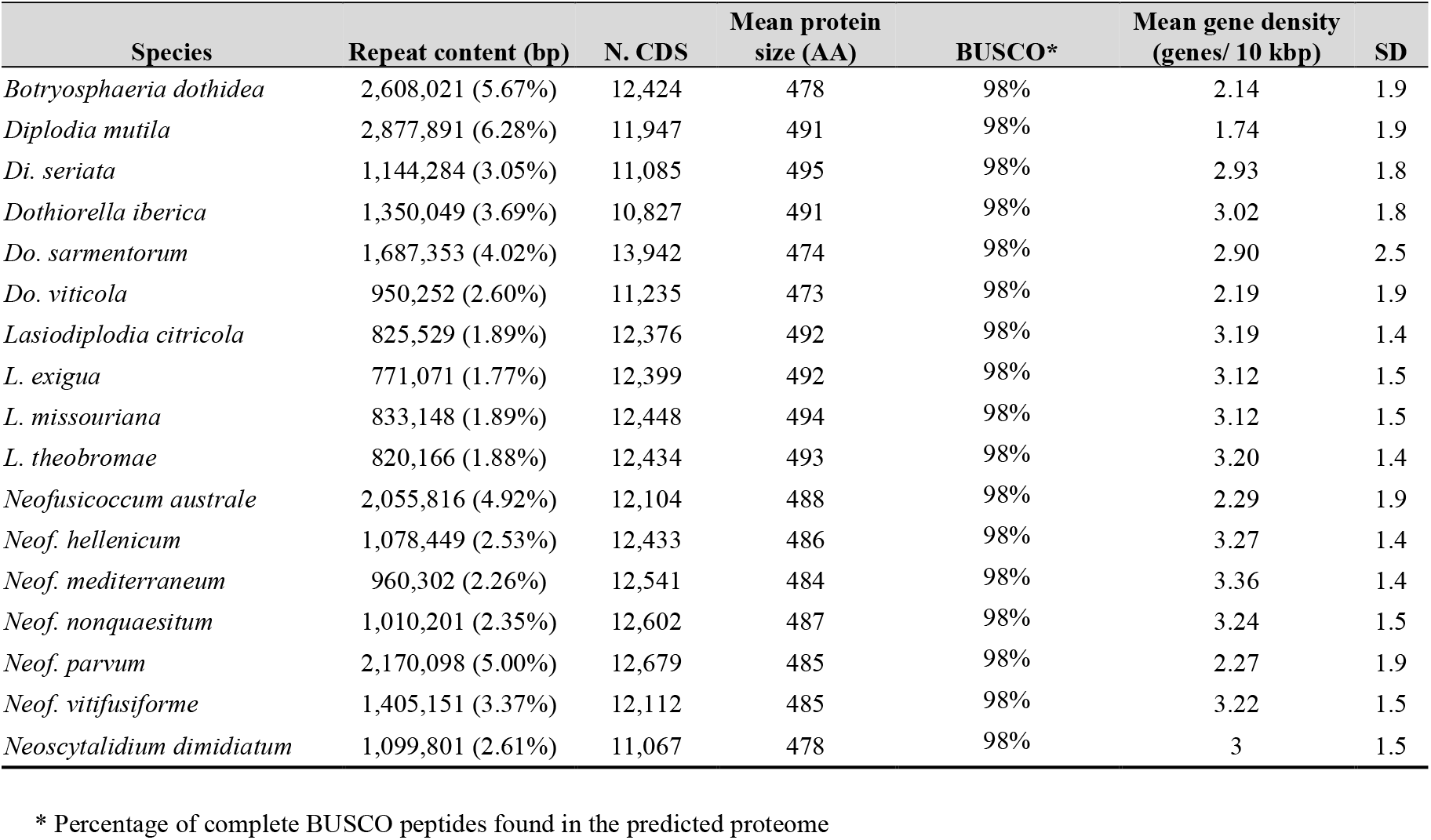
Gene model predictions statistics of the *Botryosphaeriaceae* species analyzed.

The predicted genes of the seventeen genomes were annotated using general databases for protein domains (Pfam), gene ontology (GO), as well as more specialized databases related to putative virulence factors. The last group included carbohydrate-active enzymes (CAZymes), cytochrome P450s, peroxidases, usually associated with host colonization and wood degradation, and secondary metabolism gene clusters, including toxins production and cellular transporters (**Additional File 1 Table 4**; **Table 3**). A total of 229,251 predicted protein-coding genes were annotated (**Table 3**).

**Table 3.**
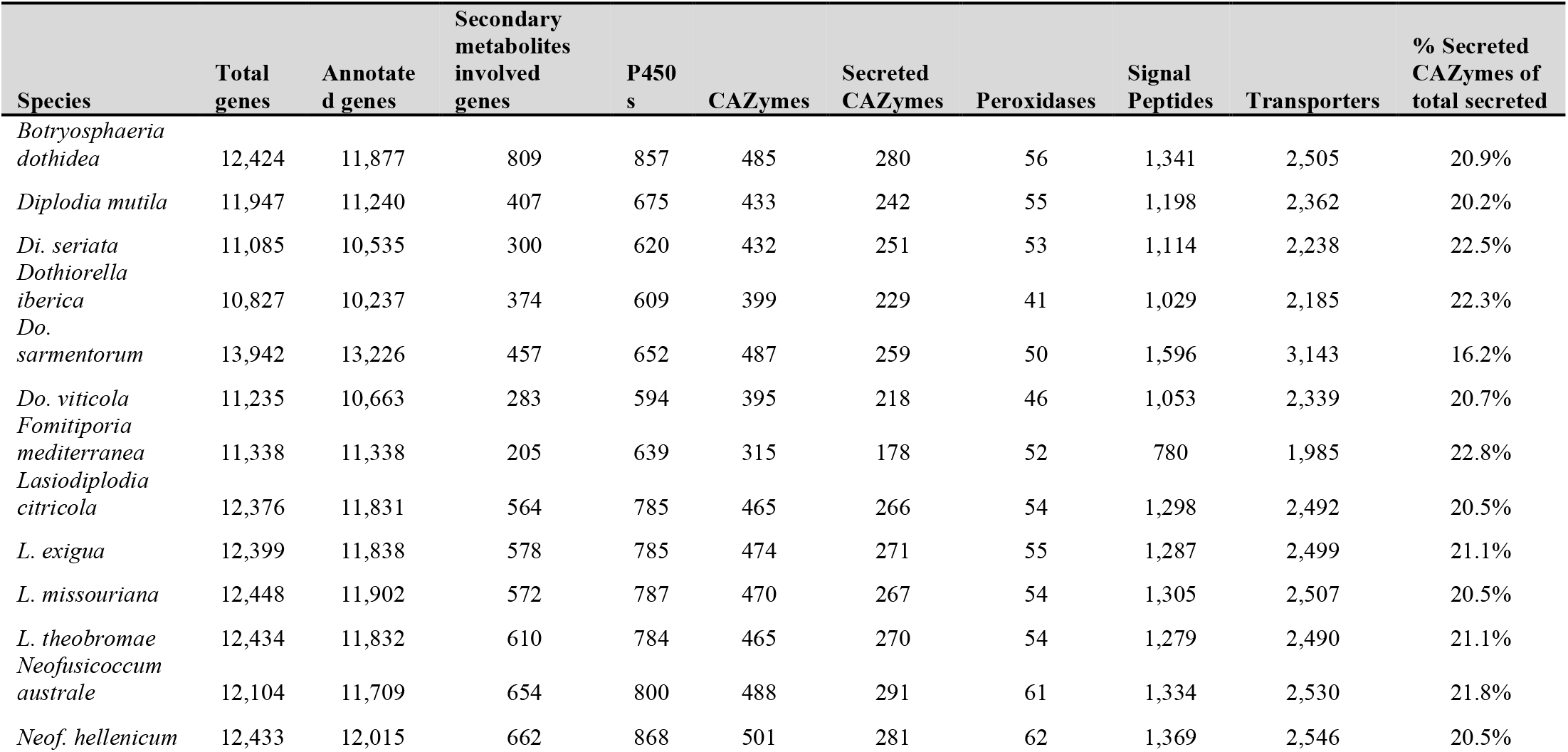
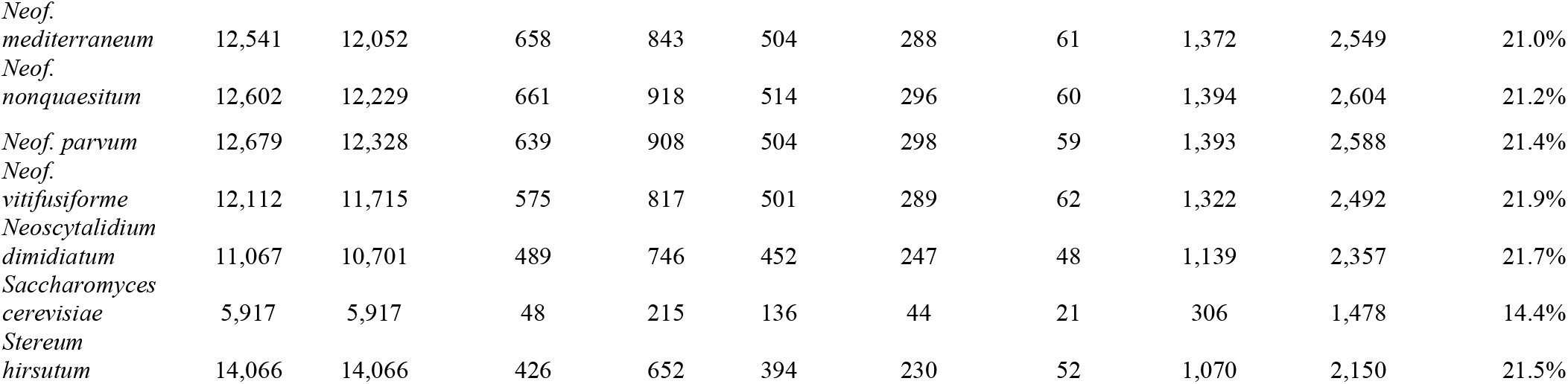
Number of protein coding genes annotated per functional category.

### Carbohydrate-active enzymes are especially abundant in the *Neofusicoccum* genus

A wide variety of monosaccharides can be linked to most types of molecules (proteins, lipids, nucleic acids and, sugar themselves) converting these glycoconjugates into one of the most structurally diverse substrates (Cantarel et al., 2009). CAZymes are the group of enzymes responsible for the assembly and breakdown of these diverse substrates (Lombard et al., 2014). Not all CAZymes contribute to the pathogenicity of the microorganisms, however predicting them in conjunction with signal peptides is widely used to obtain information about plant pathogen cell wall degrading enzymes (Morales-Cruz et al., 2015; Jones et al., 2014; Floudas et al., 2012; Suzuki et al., 2012; Blanco-Ulate et al., 2014). An average of 20.9 ± 0.3% of the predicted secreted proteins among all seventeen *Botryosphaeriaceae* genomes shared similarity with the CAZymes in the dbCAN2 database (Zhang et al., 2018). Glycoside Hydrolases (GH) and Auxiliary Activity CAZymes (AA) were the two groups with the most predicted proteins. GHs were especially abundant in *Neofusicoccum* spp. with an average of 336 ± 4 proteins compared to 303 ± 7 for the rest of the *Botryosphaeriaceae* species in this study (**Additional file 1 Table 5**). A total of 15 putative genes of GH3 were present in *Neof. nonquaesitum* and *L. missouriana*, as well as 14 in *Neof. parvum, L. citricola*, and *L. exigua*. GH3 and GH43 families activities include β-glucosidases, β-xylosidases, glucanases, L-arabinofuranosidase, galactanase and others related to the hydrolysis of plant cell wall components into more simple sugars (Sampedro et al., 2017; Cairns & Esen, 2010; Knob et al., 2010; Faure 2002; Polizeli et al., 2005).

AAs were also more abundant in *Neofusicoccum* (127 ± 3) than the rest of the *Botryosphaeriaceae* species (100 ± 4). The AA3 family was the most abundant with numerous copies in the genus *Neofusicoccum* (**Figure 2**), ranging from 21 to 26 predicted proteins in *Neofusicoccum parvum* (UCD646So). The genome of *Botryosphaeria dothidea* was predicted to possess 23 AA3 proteins. The AA3 and AA9 families include cellobiose dehydrogenases, alcohol oxidases, pyranose oxidase, acting over more complex substrates of the plant cell wall like cellulose and/or lignin (Henriksson et al., 2000; Harreither et al., 2011; Hernández-Ortega et al., 2012; Daniel et al., 1994).

**Figure 2.**
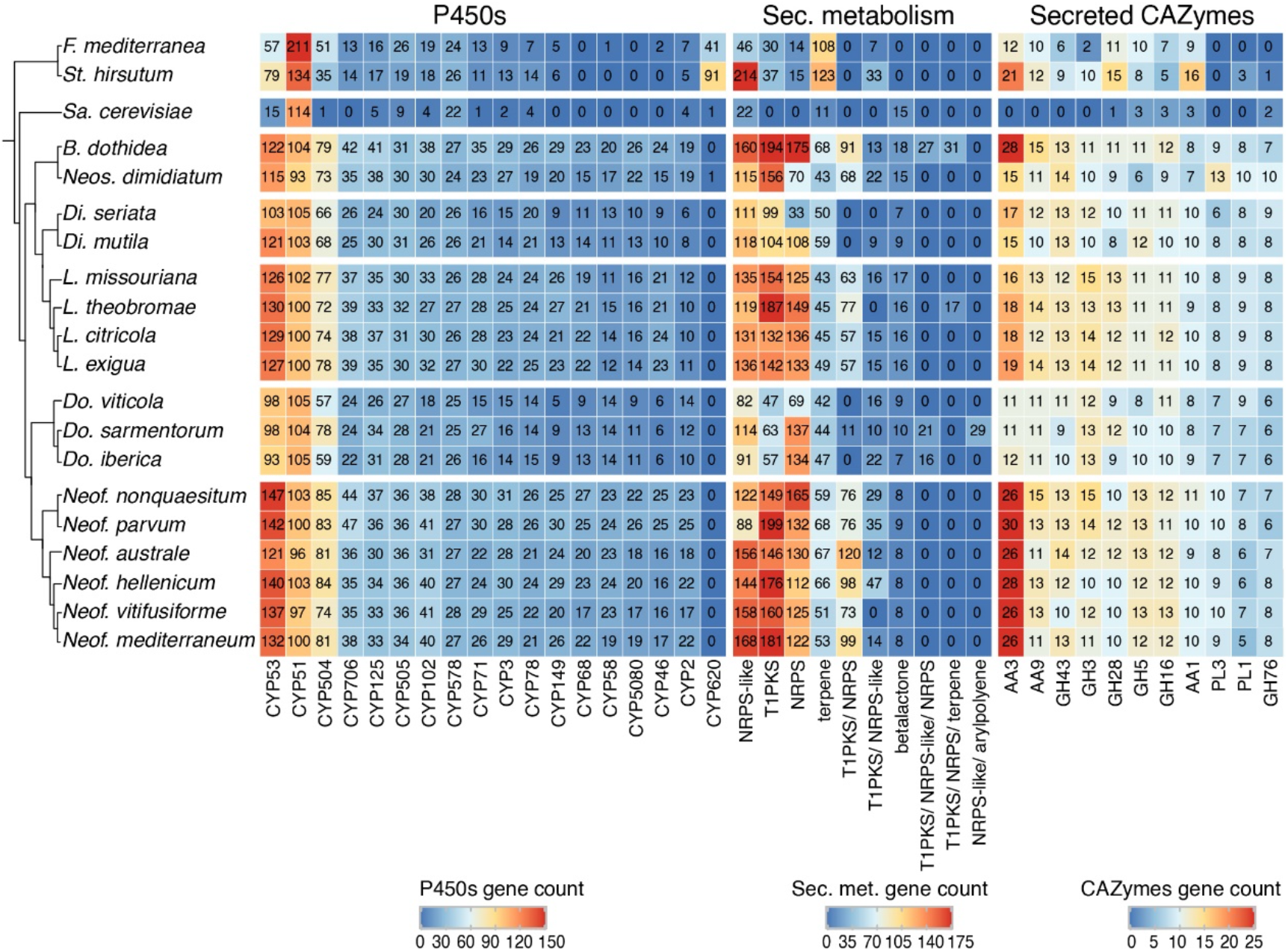
Number of protein-coding genes annotated as P450s, secondary metabolism and secreted CAZymes. The heatmap includes only the annotations with the highest number of genes across all genomes.

### *Neofusicoccum, Botryosphaeria*, and *Lasiodiplodia* species encode the largest number of predicted P450s

Cytochrome P450 enzyme evolution is thought to contribute to the adaptation of organisms to new ecological niches. The functions may vary from primary metabolism, detoxification of xenobiotic compounds, to producing a vast variety of secondary metabolites (Chen et al., 2014b; Črešnar & Petrič, 2011; Moktali et al., 2012). These features sometimes play essential roles in pathogenesis (Moktali et al., 2012; Črešnar & Petrič, 2011). The P450s were classified in superfamilies as described by Fischer et al. (2007). *Neofusicoccum, Botryosphaeria*, and *Lasiodiplodia* species encoded a larger number of predicted P450 (859 ± 20, 857, and 785 ± 1 respectively) compared to *Diplodia* and *Dothiorella* species (648 ± 28 and 618 ± 17 respectively). *Neof. nonquaesitum* and *Neof. parvum* showed the highest number of predicted P450s genes with 918 and 908, respectively (**Table 3**; **Additional file 1 Table 6**). CYP53, CYP51, and CYP504 were the most abundant across all the species. CYP53 was especially numerous in *Neofusicoccum* (137 ± 4 genes), *Lasiodiplodia* (128 ± 1 gene), and *Botryosphaeria* (119 ± 4 genes) species compared to the other genera (93±8 genes). On the other hand, the CYP51 was very consistent in the *Botryosphaeriaceae* family (from 93 to 105 genes). Other superfamilies like CYP706, CYP102, and CYP3 show the same pattern of higher representation in *Neofusicoccum, Lasiodiplodia*, and *Botryosphaeria* than in *Diplodia* and *Dothiorella* species.

### Peroxidases are most abundant in the genomes of *Neofusicoccum* species

Fungal peroxidases are oxidoreductases that use peroxides to catalyze the oxidation of various compounds ranging from ligninolysis to the detoxification of host-derived reactive oxygen species and have been shown to contribute to virulence (Choi et al., 2014; Molina, & Kahmann, 2007; Guo et al., 2010). The annotation of these peroxidases was based on the manually curated Fungal Peroxidases Database fPoxDB (Choi et al., 2014). *Neofusicoccum* species encoded the largest number of predicted peroxidases (61 ± 0), followed by *Lasiodiplodia, Botryosphaeria*, and *Diplodia* with an average of 54 ± 1 annotated genes (**Table 3**). *Dothiorella*, with only 46 ± 3 was the genus with the least number of annotated peroxidases in the *Botryosphaeriaceae* (**Additional File 1 Table 7**). Hybrid Ascorbate-Cytochrome C peroxidases were more abundant in *Neofusicoccum, Botryosphaeria*, and *Lasiodiplodia*, ranging from 8 to 11 genes, while haloperoxidases were more abundant in the genus *Neofusicoccum* (11 ± 0 genes).

### The genomes of *Botryosphaeria, Neofusicoccum*, and *Lasiodiplodia* species have the largest number of secondary metabolism gene clusters

Secondary metabolites play important roles in fungal development and interactions with other organisms, including plant hosts (Keller, 2019). Phytotoxic metabolites, e.g., melleins, produced by *Neof. parvum* both *in vitro* and in the wood of symptomatic grape are thought to be associated with pathogenesis (Abou-Mansour et al., 2015). In fungi, the genes encoding the functions responsible for the biosynthesis of secondary metabolites are physically grouped in clusters of contiguous genes (Brakhage, 2013; Keller., 2019), which typically comprise a central biosynthetic gene as well as genes involved in post-synthesis modification of the metabolites and cellular transport.

Using antiSMASH 5 (Blin et al., 2019), we detected in the seventeen *Botryosphaeriaceae* an average of 43 ± 3 biosynthetic gene clusters (BGCs). The Type I Polyketide synthase cluster (T1PKS) and the Non-ribosomal peptide synthetase-like fragment (NRPS-like) together accounted for 47% of all annotated BGCs. BGCs were most abundant in *Botryosphaeria, Neofusicoccum*, and *Lasiodiplodia* species with an average of 57 ± 8, 56 ± 1, and 49 ± 1 BGCs, respectively. In these genera, we also found the larger number of genes per BGC (11 ± 1, 12 ± 0, and 12 ± 0, respectively; **Additional file 1 Table 9**). In *Neofusicoccum* spp., 169 ± 7 genes were associated with T1PKS, 154 ± 12 in *Lasiodiplodia* and 175 ± 19 in *Botryosphaeria* (**Figure 2**). For these secondary metabolites as well as classes we found fewer genes in the genomes of *Diplodia* and *Dothiorella* species (**Additional file 1 Table 10**).

Toxins and other secondary metabolites are exported by cellular transporters (Del Sorbo et al., 2000). Homologies with the Transporters Classification Database (TCDB; Saier Jr et al., 2006) were used to annotate hypothetical protein transporters. Overall, the Electrochemical Potential-driven Transporters was the most prominent group across all the species representing 31 ± 1% of the annotated transporters followed by the Primary Active Transporters (19%) and the Incompletely Characterized Transport Systems (19%). More specifically, The Major Facilitator Superfamily (MFS) (TCDB code 2.A.1) represented the highest number in all the species but was especially abundant in *Neofusicoccum, Lasiodiplodia*, and *Botryosphaeria* from 455 to 514 predicted genes (**Additional file 1 Table 8**). The genome of *Do. sarmentorum* encodes a higher number of genes in the ATP-binding Cassette (ABC; 134 genes) superfamily compared to the other fungi analyzed (59 ± 1 genes). Both MFS and ABC transporters can be involved in toxin secretion and defense responses (Perlin et al., 2014; Del Sorbo et al., 2000).

### Estimation of gene family expansion and contraction and evaluation of functional enrichment

We further evaluated the differences in putative virulence factors content to identify gene families that have significantly expanded or contracted in specific lineages by statistical analysis of the evolution of the size of gene families using Computational Analysis of gene Family Evolution (CAFE; De Bie et al., 2006). CAFE estimates the global birth and death rate of gene families and identifies those families that have an accelerated rate of gain or loss (De Bie et al., 2006; Hahn et al., 2005). CAFE uses as input a clock-calibrated phylogenetic tree and gene family sizes in all the species’ genomes. We included in the analysis *Fomitiporia mediterranea* and *Stereum hirsutum*, two well-known wood decay basidiomycetes related to the white-rot symptom in Esca disease of grapevines, and *Saccharomyces cerevisiae*. These additional species allowed to use as calibration points the estimated dates of monophyletic partition of *Ascomycota* (588 mya) and *Dothideomycetes* (350 mya) according to Floudas et al. (2012) and Beimforde et al. (2014). To construct the phylogenetic tree, we identified twenty-one single-copy protein sequences that were previously used to study phylogenetic relationships across Fungi (Floudas et al. 2012). The phylogenetic tree was built using a multiple alignment comprising 12,066 amino acid positions. The topology of the clock-calibrated tree was confirmed independently (**Additional file 3**) using ITS (Internal Transcribed Spacer) and TEF (Translation Elongation Factor), and was consistent with published ones (Phillips et al., 2008; Thambugala et al., 2014; Chen et al., 2014a).

The gene families were computed using a Markov Cluster algorithm (MCL) that groups putative orthologs and paralogs (Enright et al., 2002). In total 237,976 proteins of the 20 fungal genomes were clustered into gene families (e-value < 1e^−6^). These family sizes and the clock calibrated tree produced by BEAST allowed CAFE to detect 666 families (35,498 genes) across all the species with a significantly higher than expected rate of gene gain/loss (*P* ≤ 0.01). The numbers of gene families expanded and contracted for each branch of the phylogeny are shown in **Figure 3**. The parent branches of the *Neofusicoccum, Lasiodiplodia*, and *Botryosphaeria* clades show a positive rate of gene gain/losses (+0.45, +0.14, and +0.44 respectively), which suggest an expansion of some set of proteins. On the other hand, the parent branches of *Diplodia, Dothiorella*, and Basidiomycetes (*Stereum* and *Fomitiporia*) clades present a negative rate of gain losses (−0.40, −0.49, and −0.70, respectively). *Saccharomyces cerevisiae*, as seen in previous studies (Morales-Cruz et al., 2015), showed the lowest rate with −1.63.

**Figure 3.**
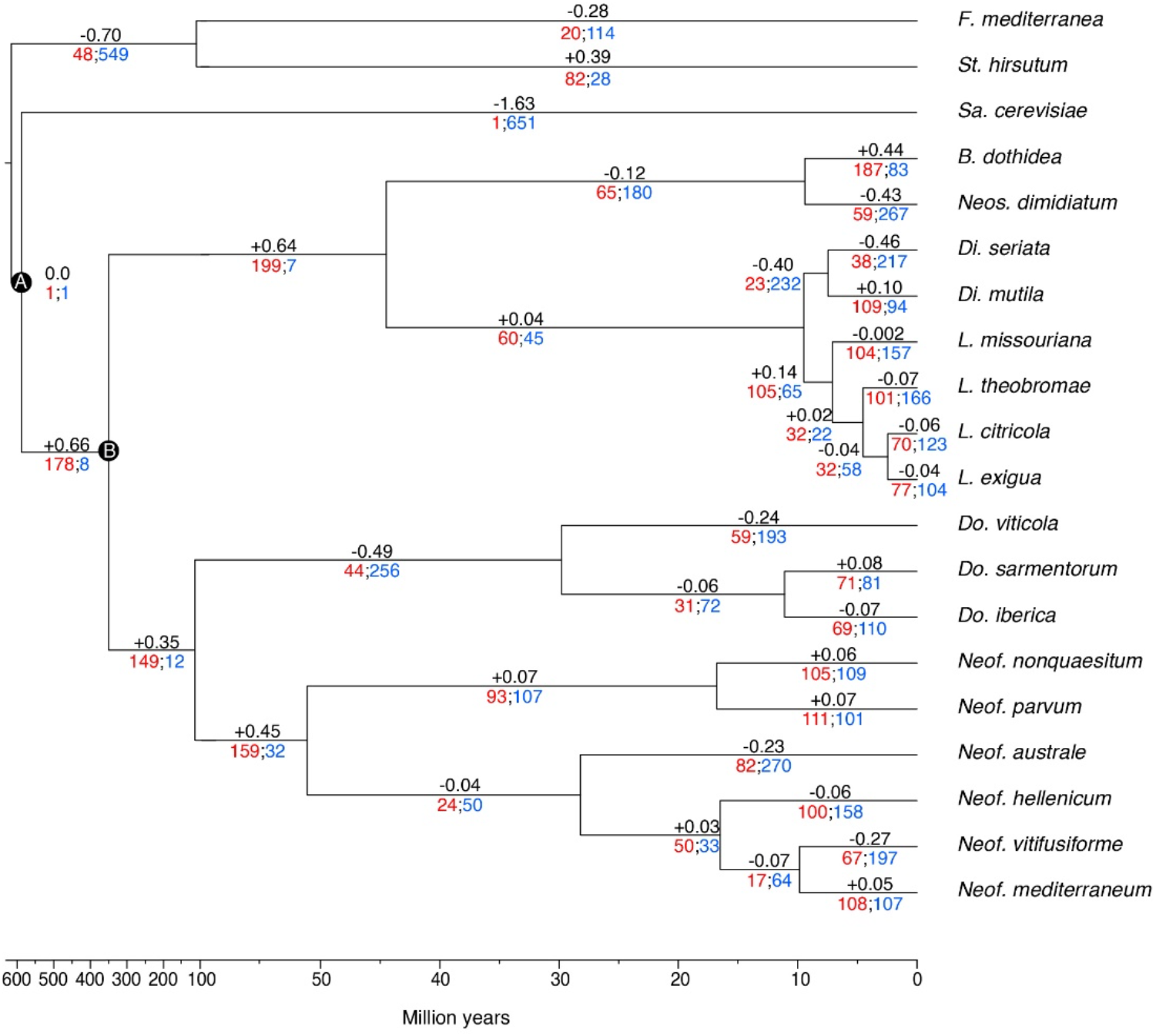
Clock calibrated phylogenetic tree showing the number of gene families significantly expanded (red), contracted (blue) and their average pattern (black).

**Figure 4.**
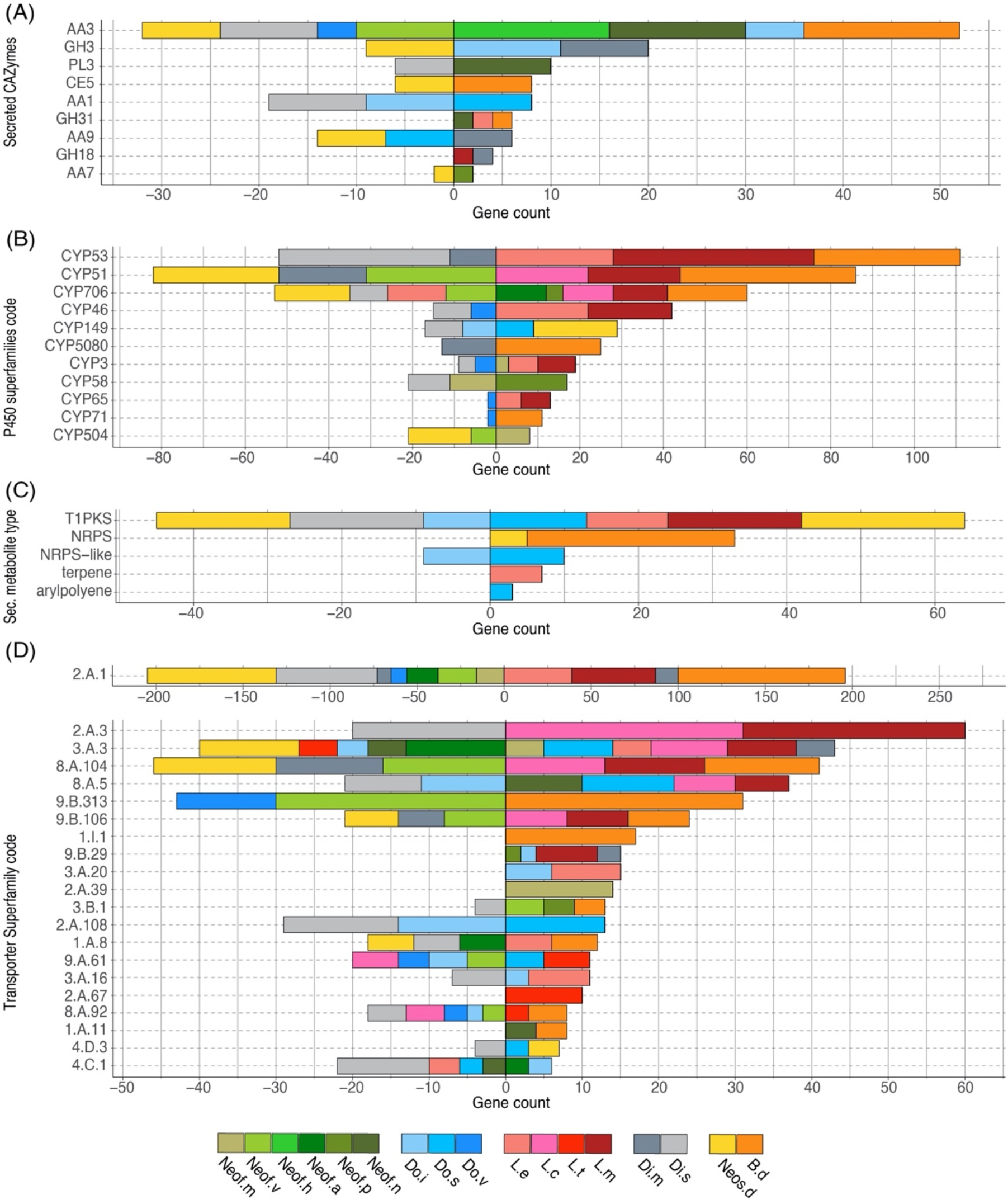
Bar plot of the counts of genes annotated in each group of significantly expanded functional category. Only genes significantly overrepresented (*P* ≤ 0.01) in the gene families expanded in the *Botryosphaeriaceae* group are shown.

The 35,498 genes in the significantly expanded or contracted families were analyzed with a Fisher’s Exact test to identify functional enrichments within those families. We found that these enrichments were not always present in all the species of a clade (**Figure 3**). However, there are some patterns to highlight. The *Lasiodiplodia* clade represents an overall expansion of the transporter proteins. *B. dothidea* also shows a significant expansion of this family. On the other hand, the *Diplodia* and *Neofusicoccum* clades show an overall contraction of transporters. The *Dothiorella* clade and *Neos. dimidiatum* shows no specific pattern. The expanded secondary metabolite proteins were specifically abundant in *L. misouriana, L. exigua, B. dothidea, Do. sarmentorum* and *Neos. dimidiatum*. The P450 family was expanded mostly in *Lasiodiplodia* species and *B. dothidea* but contracted in *Diplodia* species and *Neos. dimidiatum. Neofusicoccum* species and *B. dothidea* have an important representation of expanded Secreted CAZymes, *Diplodia* and *Dothiorella* represent several expanded proteins, however, the numbers in *Lasiodiplodia* are extremely low.

Transporter related genes in the Major Facilitator Superfamily (MFS-2.A.1) were the most enriched in all the clades analyzed (196 predicted proteins were expanded and 205 contracted). The secondary metabolite related proteins type 1 Polyketide Synthases (T1PKS) were expanded in *Neos. dimidiatum, L. missouriana, L. exigua*, and *Do. sarmentorum*, whereas Non-Ribosomal Peptide Synthetases (NRPS) were expanded in *B. dothidea* and *Neos. dimidiatum*. For the secreted CAZymes, *Neof. nonquaesitum, Neof. hellenicum* and *B. dothidea* show an enrichment of the Auxiliary Activity family 3. Also, *Do. iberica* and *Dip. mutila* show an enrichment of the Glycoside Hydrolase Family 3.

### Phylogenetically informed principal component analysis of the expanded gene families associated with virulence factors

To identify similarities between species in the *Botryosphaeriaceae* family, a phylogenetically informed-principal component analysis (phylo-PCA) was applied to the significantly expanded families of virulence functions. These gene families were grouped into the functional categories based on the specialized databases, and the PCA was carried out using the Phyl.PCA (Revell 2012). Phyl.PCA considers correlations among species due to phylogenetic relatedness, while correcting the matrices for non-independence among observations (Revell & Collar, 2009). Two separate analyses were conducted using the clock-calibrated tree presented previously and the tables of the number of genes classified as Secreted CAZymes (**Figure 5A**) and Secondary Metabolism (**Figure 5B**).

**Figure 5.**
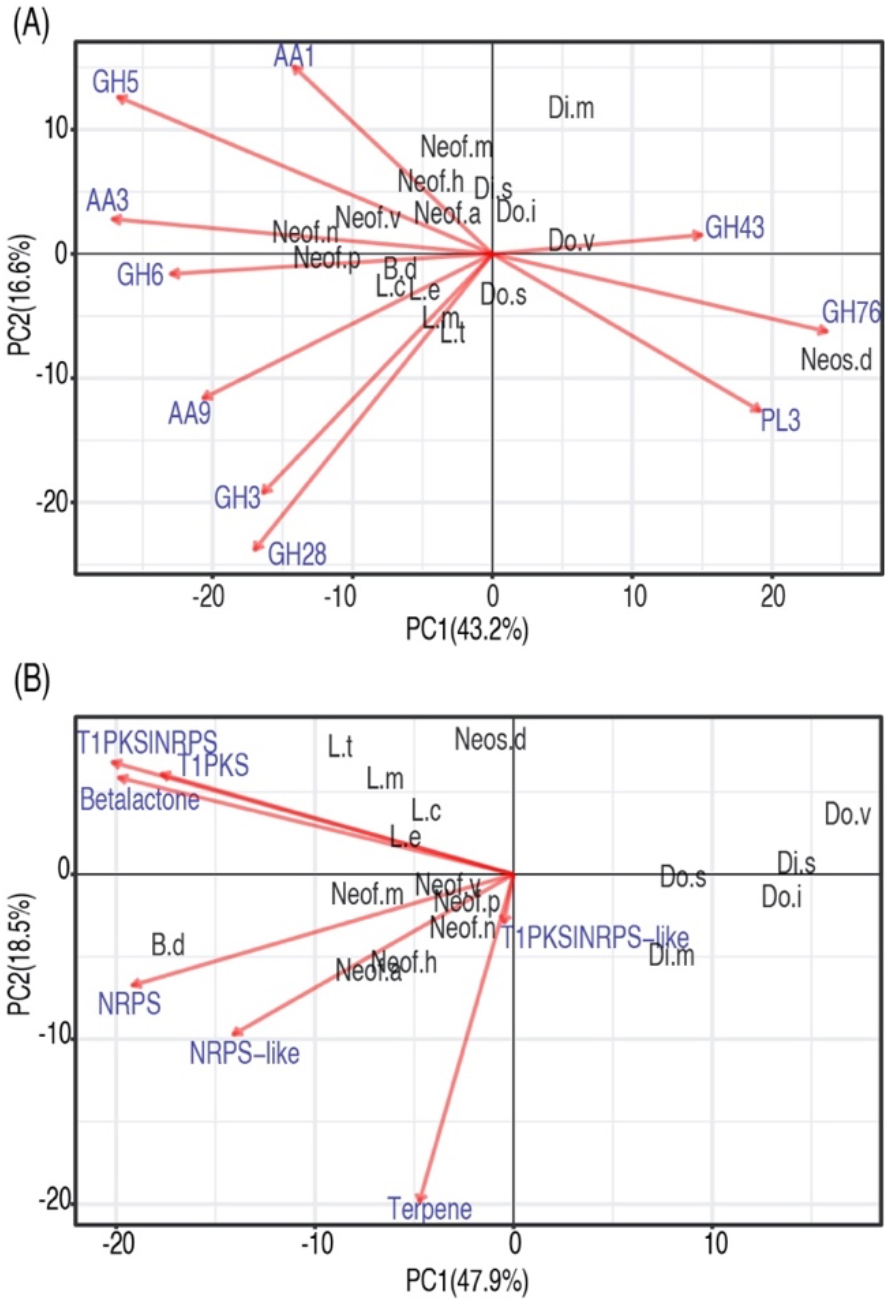
Phylogenetic principal component analysis (PCA) of the expanded gene families grouped by Secreted CAZymes (A) and Secondary metabolism clusters (B). Only vectors of the largest loadings are shown.

Due to the close phylogenetic relationship of the *Botryosphaeriaceae* family, the set of secreted CAZymes is remarkably similar. However, there is a clear separation of the species that are considered to be more virulent (**Figure 5A**), those belonging to the genera *Neofusicoccum, Lasiodiplodia*, and *Botryosphaeria* (Úrbez-Torres *et al*., 2008; Úrbez-Torres & Gubler, 2009). At the same time, we observe a close cluster of *Neofusicoccum* species which are separated from the other groups mostly by the abundance of AA1, AA3, and GH5. In addition, the genus *Lasiodiplodia* is tightly clustered together with *B. dothidea*. This is driven by the abundance of AA9, GH28, and GH3, with the last family being more abundant in *Lasiodiplodia* species. The close clustering of *Neofusicoccum, Botryosphaeria*, and *Lasiodiplodia* is driven mostly by their similar profile of GH16 and AA3. *Neos. dimidiatum* is well separated from the rest of the species by the higher presence of GH76 and PL3 proteins.

The secondary metabolism PCA shows a similar separation of the most virulent genera from the others (**Figure 5**). *Lasiodiplodia* species are grouped together by similarly high profiles of T1PKS, Betalactone and T1PKS/NRPS clusters. *Neofusicoccum* species are grouped due to high numbers of terpene synthases and NRPS-like clusters. *B. dothidea* is separated because of its high abundance of NRPS, T1PKS, Terpenes, Betalactone, and NRPS-like clusters.

## Discussion

In this study, we describe the genome sequences of seventeen well-known canker-causing fungal species in the *Botryosphaeriaceae*. The genomes assembled coupled with *in-planta* experiments allowed us to start analyzing the pathogenicity levels and the virulence factor profiles within this important fungal family. The level of completeness of the assembled genomes is consistent across all the drafts based on the expected and assembled genome sizes. This behavior is also confirmed by the high representation of conserved genes (Simão *et al*., 2015; Parra *et al*., 2007). The completeness of the genomes, as well as the protein-coding genes and the repetitive DNA content, are similar to those of other wood-colonizing fungi of grape, such as *Diaporthe ampelina DA912* (Morales-Cruz et al., 2015), *Diplodia seriata DS831* (Morales-Cruz et al., 2015), and *Lasiodiplodia theobromae LA-SOL3* (Félix et al., 2019). Apart from the estimated completeness of the genomes, it is necessary to understand some of the limitations of the short reads technology, like copy number errors, chimeric contigs, and under-representation of repetitive regions (Treangen & Salzberg, 2012, Alkan et al., 2011).

The functional annotation of the seventeen *Botryosphaeriaceae* species presents a broad and variable profile of virulence factors that are used in different ways by fungi to colonize and survive in their hosts (Schulze-Lefert & Panstruga, 2011; Peyraud et al., 2019). The results show a great variation in the number of genes identified with a functional category, and these differences were usually associated with the genus of each species like those observed by Baroncelli et al. (2016) in *Colletotrichum* and Morales-Cruz et al. (2015) in other grapevine trunk pathogens. Researchers are inclined to think that the gene content is associated with the lifestyle and the variety of hosts (Baroncelli et al., 2016; Zhao et al., 2013; Lo Presti *et al*., 2015). The expansion or contraction of a gene family usually occurs on functions that are under positive or negative selection. For instance, the genes related to host colonization and defense are under high pressure, therefore, it is common to encounter duplications or even losses. On the other hand, genes related to growth are more conserved and usually are selected against these changes (Wapinski *et al*., 2007). Gene duplication events are crucial as they are considered to be one of the main processes that generate functional innovation (Ohno, 2013; Zhang 2003). This process plays one of the most important roles in fungal adaptation and divergence (Gladieux *et al*., 2014).

Host colonization during infection is mostly driven by gene expression of some groups of well-known proteins, namely, the secreted CAZymes, cytochrome P450 monooxygenases, peroxidases, and secondary metabolite-producing proteins (Massonnet *et al*., 2018). The *Botryosphaeriaceae* family has a variable profile of these sets of genes, with the most virulent and aggressive species having, on average, greater numbers of annotated genes in these categories (**Table 3**). In grape and pistachio, species in the genera *Neofusicoccum* and *Lasiodiplodia*, are typically more virulent than species in the genera *Diplodia* and *Dothiorella* (Nouri et al., 2019; Úrbez-Torres *et al*., 2008; Úrbez-Torres & Gubler, 2009). GH functions of β-glucosidases, β-xylosidases, glucanases, L-arabinofuranosidase, and galactanase were present in all the pathogens in this study and significantly more in *Neofusicoccum* and *Lasiodiplodia* . In the same way as the GH, AA functions like cellobiose dehydrogenases, alcohol oxidases, pyranose oxidase were more abundant among *Neofusicoccum* species and *Botryosphaeria dothidea*. GH and AA play a critical role in the degradation of the host cell wall compounds (Kubicek *et al*., 2014), which is involved with the degree of pathogenicity within these genera, albeit on grape, the host we examined. Massonnet et al. (2018), Félix et al. (2019) and Marsberg et al. (2017), found similar numbers of CAZymes in *Neof. parvum, L. theobromae*, and *B. dothidea*, respectively. P450s are instrumental to the development of all organisms. These enzymes are involved in many aspects of primary and secondary metabolisms and are responsible for xenobiotic detoxification and degradation (Črešnar & Petrič, 2011; Moktali et al., 2012). Virulence may in part reflect the ability of some species to better tolerate and, further, to metabolize phenolic compounds produced by the host. Both *N. parvum* and *D. seriata* can eliminate the stilbene piceid and its derivative resveratrol *in vitro* (Stempien et al. 2017), but the former is better able to tolerate resveratrol derivatives ampelopsin A, hopeaphenol, isohopeaphenol, miyabenol C, and ε-viniferin, which are produced at higher levels *in planta* in response to *N. parvum* versus *D. seriata* infection (Lambert et al., 2012). Therefore, it is not unexpected to see a variable profile amongst genera in the *Botryosphaeriaceae* family and even within a single genus. As presented in **Figure 2**, some superfamilies are abundant in *Neofusicoccum, Lasiodiplodia* and *Botryosphaeria* genera, but other superfamilies are especially more numerous in the Basidiomycetes species included in this study. On the other hand, for most of the superfamilies presented, *Sa. cerevisiae* shows a considerable lack of such annotated genes, but CYP53 and CYP578 the counts are comparable with the rest of the species. This variation is sourced by the constant evolution and adaptation of the microorganism and hosts to their specific environment (Yan *et al*., 2018).

As plants evolve new defense mechanisms and compounds against pathogens, the fungi diversify their methods to degrade these compounds or generate new metabolites to attack their hosts (Yan *et al*., 2018; Deng *et al*., 2007). The *Botryosphaeriaceae* species in this study and the two Basidiomycetes present a set of fungal peroxidases that range from 41 to 62. As for the previous putative virulence factors, *Neofusicoccum, Lasiodiplodia* and *Botryosphaeria* genera have the most annotated peroxidases, however in this case, *Diplodia* also showed a comparable amount. Manganese peroxidase was only found in the two basidiomycetes. This enzyme has a critical role in the degradation of lignocellulose compounds by basidiomcyetes (Elisashvili & Kachlishvili, 2009; Liers *et al*., 2011), therefore it is very common in white-rot fungi such as *F. mediterraneum* and *St. hirsutum* (Morgenstern *et al*., 2010, Lee *et al*., 2015). Ascomycetes that rot wood are characterized as soft-rot fungi, which do not degrade lignin by producing manganese peroxidase, but instead ‘alter’ lignin (to gain access to cellulose and hemicellulose) by producing lignin peroxidases, peroxidases, polyphenol oxidases, and laccases (Goodell et al. 2008). Haloperoxidases also have roles in lignin degradation and toxic compound resistance (Hofrichter *et al*., 2010: Mayer *et al*., 2001; Zámocký & Obinger, 2010). The former enzyme was found in higher numbers in the genus *Neofusicoccum* compared to other genera within the family. The hybrid ascorbate-cytochrome C peroxidase was overrepresented in the genera *Neofusicoccum, Lasiodiplodia* and *Botryosphaeria* and is associated directly with the detoxification of ROS (Wang *et al*., 2016; Zámocký *et al*., 2014; Segal & Wilson, 2018).

The wide array of transporters annotated in this study suggests a high adaptation to toxic compounds, either produced by other microorganisms, the host, or potentially chemical synthesized fungicides (Stergiopoulos *et al*., 2002). The number of proteins in the Major Facilitator Superfamily (MFS) and Superfamily in *Neofusicoccum, Lasiodiplodia* and *Botryosphaeria* were more numerous than the other *Botryosphaeriaceae* species. Protein members of the MFS family may have different functions in the influx/efflux of molecules between cells and the exterior environment, and several cases of fungicide resistances have been associated with the overexpression of certain MFS channels (Stergiopoulos *et al*., 2002; Chen *et al*., 2017; Dos Santos *et al*., 2014; Gulshan & Moye-Rowley, 2007). The former genera have been reported to have lower sensitivities to almost full resistance to different synthetic fungicides (Tennakoon *et al*., 2019; Li *et al*., 2020; Wang *et al*., 2010). Similar behavior was observed in *Dothiorella sarmentorum*, were the ATP-binding Cassette (ABC) is highly represented. The ABC superfamily plays different roles in fungicide resistance, mycelial growth and overall pathogenicity (Stergiopoulos *et al*., 2002; Qi *et al*., 2018). In addition, the array of secondary metabolite gene clusters is more expanded in the *Botryosphaeriaceae* family than in the Basidiomycetes except for terpene synthase gene clusters. T1PKS, NRPS, and hybrids of T1PKS-NRPS produce toxic polyketides and toxic polypeptides, which kill host cells and leads to disease development (Blin et al., 2019; Andolfi *et al*., 2011; Pusztahelyi *et al*., 2015; Luini *et al*., 2010; Morales-Cruz et al., 2015).

To evaluate with more detail the potential differences in virulence within the *Botryosphaeriaceae* family, we executed a Computational Analysis of gene Family Evolution (De Bie et al., 2006). By identifying species and gene families with higher rates of gain and loss can help us to better understand the differences in pathogenicity as it relates to the numbers of copies of virulence genes (Morales-Cruz et al., 2015; Hahn et al., 2005). Six hundred and sixty-six gene families of the proteins analyzed in this study have a significantly higher than expected rate of gain/loss. The annotation of putative virulence factors in *Neofusicoccum, Lasiodiplodia*, and *Botryosphaeria* shows an average expansion of these gene families, even if some of the species shows a contraction, the overall clade rate is positive. Among those expanded or contracted families there is a set of functions that are overrepresented. The secreted CAZymes seem to be expanding in *Neof. hellenicum, Neof. nonquaesitum, B. dothidea, Di. mutila, Do. iberica*, and *Do. sarmentorum*, whereas the *Dothiorella* species show contractions in some families. However, almost no significant gain/loss of secreted CAZymes appears to be occurring in the genomes of *Lasiodiplodia* species. The opposite scenario is observed for the P450s, where *Lasiodiplodia* appears to be actively evolving, showing major expansions in three of the four species in this study. Also, *B. dothidea* and three *Neofusicoccum* species (*Neof. parvum, Neof. australe* and *Neof. mediterraneum*) show an expansion of these families. On the other side, *Neos. dimidiatum, B. dothidea, Do. sarmentorum, L. exigua*, and *L. missouriana* are actively expanding their secondary metabolite gene clusters. Finally, the wide variety of transporters present in fungi, is the result of the positive selection pressure over them, the need of the fungi to adapt to new environments and hosts had selected for multiple mutations that diversifies the transporters functions (Gladieux *et al*., 2014). The MFS (2.A.1) displays the largest effect of expansion and contraction among all the species. *B. dothidea, L. missouriana, L. exigua*, and *Di. mutila* appear to be actively expanding the MFS transporters. However, *Neos. dimidiatum, Di. seriata, Neof. vitifusiforme, Neof. australe* and *Neof. mediterraneum* are contracting MFS transporters.

Phylo PCAs results support the idea that within the *Botryosphaeriaceae* family, *Neofusicoccum, Lasiodiplodia*, and *Botryosphaeria* genera are the most virulent (Úrbez-Torres *et al*., 2008; Úrbez-Torres & Gubler, 2009). There was a very clear separation of these species from the *Diplodia, Dothiorella* and *Neoscytalidium*. The secreted CAZymes that cause the clustering of the *Neofusicoccum* species are usually associated with laccases, cellobiose dehydrogenases, and cellulase activities. These enzymes usually target components of the plant cell wall such as lignin, cellulose, cellobiose (Fillat *et al*., 2016; Cameron & Aust, 2001; Zamocky *et al*., 2006; Di Francesco *et al*., 2020). Among the functions driving the clustering of *Lasiodiplodia* and *Botryosphaeria*, the lytic polysaccharide monooxygenases (LPMOs, AA9) are one of the most important. They have a role in the oxidative degradation of various biopolymers such as cellulose and chitin. LPMOs can increase the activity of cellulases highly, and now, they are used in a mixture for the preparation of biofuels (Labourel *et al*., 2020, Frommhagen *et al*., 2018). Therefore, this set of enzymes may facilitate the colonization and infection of their hosts. The separation of *Neos. dimidiatum* from the rest of species is caused by GH78 which includes mannanases, α-glucosidase enzymes and the PL3 family of pectate lyases. As *Neos. dimidiatum* is also known for infecting the fruits and soft tissues of their hosts (Marques *et al*., 2013, Nouri *et al*., 2018), this set of enzymes seems to be well developed.

*Lasiodiplodia* species have a wide array of secondary metabolites. Their profile varies according to the species, isolate, and even the host (Salvatore *et al*., 2020). These metabolites are often synthesized by clusters of T1PKS, T1PKS/NRPS, and some betalactones (Salvatore *et al*., 2020, Félix et al., 2019), which are some of the major drivers for their clustering in the phylo-PCA (Figure 5) . *Neofusicoccum*, besides the previous gene clusters, also have lamanypenes and NRPS, which drives their clustering in the PCA. Similar results have been presented by Morales-Cruz et al. (2015) and Massonnet et al. (2018). Very little literature is available about the effect of the secondary metabolites of *B. dothidea* on their plant host; however, this fungus is known for its remarkable ability to produce secondary metabolites *in vitro* (Wang *et al*., 2018), which recently have been studied for their potential use as commercial antioxidants (Druzian *et al*., 2020; Valente *et al*., 2020; Xiao *et al*., 2014).

Few of the fungi in this study have been characterized in terms of their interactions with wood and individual wood components, their activation of cell-wall degrading enzymes, or their ability to tolerate phenolic compounds. Therefore, it is difficult to connect the pattern of gene family evolution to such aspects of fungal biology, especially in a comparative way among so many species, none of which have all been compared at once on a single host. The pathogenicity test on young, rooted grapevine plants raises some interesting observations. First, *L. theobromae, Neof. parvum* and *Neof. australe* are among the species that induced the most prominent lesions in the plants. These results are consistent with those of Úrbez-Torres & Gubler (2009), who found these species to be highly virulent on grape. This same study found *Di. mutila, Di. seriata, Do. iberica*, and *Do. viticola* to be weakly virulent, which is congruent with the results presented by this study. Although most *Neofusicoccum s*pecies present high numbers of putative virulence factors, the targets for these may be variable within the genus. In this pathogenicity experiment, the isolates of *Neof. vitifusiforme, Neof. nonquaesitum* and *Neof. hellenicum* were isolated from active cankers in walnut and pistachio trees, and even if some of these species can develop disease in grapevine, their virulence on *Vitis vinifera* may not be the same. Finally, the reasons why lesions produced by *B. dothidea* were not significantly different from the control are difficult to assess with certainty, but studies had reported *B. dothidea* to be weakly or moderately pathogenic on grapevine (Úrbez-Torres & Gubler, 2009; Pitt e*t al*., 2013). Some researchers remark that in potted plants after 5-6 weeks of inoculation with *B. dothidea*, the plants were not different compared to the control, whereas the other species in that study showed poor bud development and stunted green shoot growth (Úrbez-Torres & Gubler, 2009). Also, in other studies, this species is presented as endophytic and latent pathogen and they suggest that the environmental conditions can have a significant effect on the development of the disease (Piškur & Jurc, 2011; Marsberg e*t al*., 2017).

## Author Contributions

RT, KB, DPL, and DC conceived the study. DPL and RT carried out the pathogenicity experiments. PER and RHM contributed fungal strains. JG, AMC, AM, DPL, and DC carried out the computational analysis. JG, DPL, KB, and DC wrote the manuscript. All authors read and approved the final manuscript.

## Funding

This work was supported by the USDA, National Institute of Food and Agriculture, Specialty Crop Research Initiative (grant 2012-51181-19954) and by the American Vineyard Foundation (grant 1798). DC was also partially supported by the Louis P. Martini Endowment in Viticulture and the UC Davis Chancellor Fellow Award.

## Conflict of Interest Statement

The authors declare that the research was conducted in the absence of any commercial or financial relationships that could be construed as a potential conflict of interest.

## Supplementary Material

The Supplementary Material for this article can be found online at [TBD].

## Notes

### Competing Interest Statement

The authors have declared no competing interest.

